# Gene expression patterns in synchronized islet populations

**DOI:** 10.1101/377317

**Authors:** Nikita Mukhitov, Michael G. Roper

## Abstract

In vivo levels of insulin are oscillatory with a period of ~5-10 minutes, implying that the numerous islets of Langerhans within the pancreas are synchronized. While the synchronizing factors are still under investigation, one result of this behavior is expected to be coordinated intracellular [Ca^2+^] ([Ca^2+^]_i_) oscillations throughout the islet population. The role that coordinated [Ca^2+^]_i_ oscillations have on controlling gene expression within pancreatic islets was examined by comparing gene expression levels in islets that were synchronized using a low amplitude glucose wave and an unsynchronized population. The [Ca^2+^]_i_ oscillations in the synchronized population were homogeneous and had a significantly lower drift in their oscillation period as compared to unsynchronized islets. This reduced drift in the synchronized population was verified by comparing the drift of *in vivo* and *in vitro* profiles from published reports. Microarray profiling indicated a number of Ca^2+^-dependent genes were differentially regulated between the two islet populations. Gene set enrichment analysis revealed that the synchronized population had reduced expression of gene sets related to protein translation, protein turnover, energy expenditure, and insulin synthesis, while those that were related to maintenance of cell morphology were increased. It is speculated that these gene expression patterns in the synchronized islets results in a more efficient utilization of intra-cellular resources and response to environmental changes.

## Introduction

Pancreatic islets are key modulators of glucose homeostasis through the release of insulin and other hormones.^1^ Secretion of these hormones is dynamic occurring in a multi-oscillatory secretion rhythm that encompasses timescales from minutes to hours. While circadian^2^ and ultradian^3^ insulin oscillations are well understood, the rapid pulses observed *in vivo* with periods ranging from 5-to 10-min,^4-7^ are less characterized. However, it has been shown that diminished insulin oscillations are one of the first observable features of type 2 diabetes,^8^ further emphasizing the significance of the dynamic profile of hormone release.

The pulsatile nature of insulin secretion originates from the smallest structural unit within the islets, the pancreatic beta cell. Within beta cells, increases in the ratio of ATP/ADP due to increased flux through glycolysis and oxidative phosphorylation closes ATP-sensitive potassium channels with a concomitant depolarization in the cell membrane potential.^9,10^ Depolarization opens L-type Ca^2+^ channels resulting in an influx of extracellular Ca^2+^ and a rise in intracellular [Ca^2+^] ([Ca^2+^]_i_). This increase in [Ca^2+^]_i_ initiates insulin secretion through Ca^2+^-dependent processes.^11^ It is widely believed that ~5-min oscillations in glucose metabolism results in the oscillatory dynamics of the downstream processes mentioned above, producing oscillations in membrane potential,^9,10^ [Ca^2+^]_i_ [9-12], and insulin release^12,13^. The individual beta cells within an islet are synchronized by electrical coupling and diffusion of glycolytic intermediates via gap-junctions^9,10^. An elusive question that remains is how do the ~1,000,000 islets within an individual synchronize to produce a pulsatile output of hormones from the pancreas?

One long standing hypothesis is that islet synchronization occurs through classic insulin-glucose feedback loops that result in oscillatory glucose and insulin levels, both of which have been observed *in* vivo.^14-17^ This feedback loop has been mimicked in a microfluidic system by delivering glucose levels to a population of islets and iteratively adjusting the glucose levels by use of a mathematical model that mimicked insulin-dependent glucose uptake.^16,17^ With the appropriate parameters of the model, the islets synchronized resulting in population level oscillations of [Ca^2+^]_i_, insulin secretion, and glucose levels. The average period to which the system converged to was ~5 min, similar to the period of insulin secretion observed *in* vivo.^16,18^ To simplify this feedback model, these parameters can be mimicked by delivering a 5-min glucose wave to an islet population.^8, 18-20^ These “open-loop” experiments result in [Ca^2+^]_i_ and insulin oscillations that are phase-locked, or entrained, to the glucose wave.

In this study, we set out to determine if there are advantages to islet synchronization. We hypothesized that a synchronized islet population would have a more stable [Ca^2+^]_i_ oscillation period than a non-synchronized population, due to the control offered by a feedback system. These stable [Ca^2+^]_i_ oscillation periods may induce or repress specific genes in the islets. It has been shown in multiple cells types that the frequency, amplitude, and phase of [Ca^2+^]_i_ oscillations offers specificity and robustness to signal transduction pathways.^21-26^ Some of the functions encoded by [Ca^2+^]_i_ signatures include cellular proliferation and differentiation, morphogenesis, contraction, and secretion.^23^ Moreover, expression levels of several genes were found to be dependent on the oscillation frequency of [Ca^2+^]_i_.^24,26^ In one study performed with T-cells, oscillations in [Ca^2+^]_i_ activated NFAT and NF-kB more efficiently than a constant [Ca^2+^]_i_ level.^27,28^ In another study with B-lymphocytes, the activation of NFAT was highest when only select values of [Ca^2+^]_i_ oscillation frequencies were present.^29^ Finally, it has been shown that intra-portal delivery of insulin pulses induced specific gene expression in the liver as compared to the same dose of insulin delivered in a constant manner.^30^ While these effects of dynamic signals have been shown in other cell types, they have been largely ignored in pancreatic islets.

To test the hypothesis that islet synchronization may produce advantages over non-synchronized populations, the experimental mimic of the classic insulin-glucose feedback loop was performed whereby a population of islets was synchronized by exposure to a glucose wave, lysed, and their gene expression levels compared to a group of islets that were not synchronized. Between the two populations, differential regulation of Ca^2+^-dependent genes and gene sets were observed, which we hypothesize may have a positive influence on cell function in the synchronized state.

## Materials and Methods

### Chemical reagents and supplies

Cosmic calf serum, dimethyl sulfoxide (DMSO), fluorescein, 4-(2-hydroxyethyl)-1-piperazineethanesulfonic acid (HEPES), MgCl_2_, NaCl, pluronic F-127, RPMI 1640, and an antibiotic/antimycotic solution were obtained from Sigma-Aldrich (St. Louis, MO). Dextrose was obtained from Fisher Scientific (Pittsburgh, PA). CaCl_2_, KCl, and NaOH were obtained from EMD Chemicals (Gibbstown, NJ). Fura-2 acetoxymethyl ester (Fura-2 AM) was obtained from Invitrogen (Carlsbad, CA). Poly(dimethyl siloxane) (PDMS) elastomer kit was obtained from Dow Corning (Midland, MI).

For all islet stimulation experiments, a balanced salt solution (BSS) was used. BSS was composed of 125 mM NaCl, 2.4 mM CaCl_2_, 1.2 mM MgCl_2_, 5.9 mM KCl, 25 mM HEPES, and adjusted to pH 7.4. The BSS was further supplemented with either 3 or 13 mM glucose. Fura 2-AM stock was prepared by reconstitution in 10 μL of pluronic F127 and 10 μL of DMSO. The prepared stock was stored protected from light at room temperature.

### Isolation and handling of islets of Langerhans

All experiments were performed under guidelines approved by the Florida State University Animal Care and Use Committee (ACUC) protocol #1519. Islets were isolated from 20-40 g male CD-1 mice (Charles River Laboratories, Wilmington, MA) as previously described.^19,20^ All efforts were made to minimize suffering. For each set of experiments, islets were isolated from at least three mice and equally pooled from each animal (e.g., 40 islets were taken from three mice to a total of 120). From this pooled collection, islets were randomly selected for stimulation and imaging of [Ca^2+^]_i_. All experiments were conducted within two days of isolation.

### Experimental Setup

Microfluidic devices were made from PDMS and fabricated using conventional soft lithography.^31^ The design used in this study was described in previous work.^17^ Imaging of [Ca^2+^]_i_ was performed as previously described.^17^ Briefly, the microfluidic device was positioned on the stage of a Nikon Ti-S microscope equipped with a 10X, 0.5 NA objective (Nikon Instruments, Melville, NY). Excitation was achieved with a Xenon arc lamp equipped with an integrated shutter and filter wheel (Sutter Instruments, Novato, CA). Acquisition was performed on a charge coupled device (CCD) camera (Photometrics, Tucson, AZ). Nikon NIS Elements software was used to control the camera, shutter, and filter wheel. During imaging, excitation was performed sequentially at 340 +/- 5.5 and 380 +/- 5.5 nm (Chroma, Bellows Falls, VT). Emission was filtered through a 415 nm long pass dichroic mirror and a 510 +/- 20 nm emission filter (Omega, Brattleboro, VT). The response of islets is given as a ratio of fluorescence emission generated from excitation at 340 and 380 nm (F340/F380). Imaging was performed every 30-s with 100-ms exposure per excitation channel.

### Islet stimulation and imaging

Fura 2-AM was loaded into islets in serum free RPMI 1640 medium for 42 min in an incubator controlled at 37°C and 5% CO_2_. After loading with dye, 6 or 7 islets were placed in the microfluidic device and perfused with BSS containing 3 mM glucose for 1-min to wash them free of media. The experiment then commenced by perfusing islets with BSS at containing 3 mM glucose for 2-min. After 2-min, the glucose challenge was applied in either a constant manner at 11 mM or as a sinusoidal wave with a median value of 11 mM, an amplitude of 1 mM, and a 5-min period. The stimulation was maintained for 80-min.

Following the 82-min experiment, the islets were extracted and lysed using the Qiagen RNeasy plus micro kit (Qiagen, Hilden, Germany). The lysate was homogenized and held at room temperature for 10-min. The lysate was then flash frozen in liquid N2 and held at −80°C. Once all islet experiments were completed and the lysates frozen, the samples were thawed and the total RNA from each sample was purified using spin columns from the RNeasy plus micro kit. Purified total RNA was reconstituted in RNAse-free water. Approximate RNA yields were 15 ng/islet and RIN values of 10.0 were observed for all samples analyzed. Each glucose stimulation protocol was performed two more times and the total RNA from 21 islets were pooled and analyzed on a single microarray. Islets from different mice were then stimulated using one of the two glucose protocols, lysed, and pooled in the same manner for biological replicates on additional microarrays. Three Affymetrix Mouse Gene 2.0 ST microarrays were used for each glucose experimental protocol.

### Data analysis

Mouse Gene 2.0 ST CEL files were normalized to produce gene-level expression values using the implementation of the Robust Multiarray Average (RMA)^32^ in the affy package (version 1.36.1)^33^ included in the Bioconductor software suite (version 2.12)^34^ and an Entrez Gene-specific probeset mapping (17.0.0) from the Molecular and Behavioral Neuroscience Institute (Brainarray) at the University of Michigan.^35,36^ Array quality was assessed by computing Relative Log Expression (RLE) and Normalized Unscaled Standard Error (NUSE) using the affyPLM package (version 1.34.0).^37^ Principal Component Analysis (PCA) was performed using the prcomp R function with expression values that had been normalized across all samples to a mean of zero and a standard deviation of one. Differential expression was assessed using the moderated (empirical Bayesian) t-test implemented in the limma package (version 3.14.4). Correction for multiple hypothesis testing was accomplished using the Benjamini-Hochberg false discovery rate (FDR).^38^ Human homologs of mouse genes were identified using HomoloGene (version 68).^39^ All microarray analyses were performed using the R environment for statistical computing (version 2.15.1).

Gene Set Enrichment Analysis (GSEA, version 2.2.1)^40^ was used to identify biological terms, pathways and processes that were coordinately up‐ or down-regulated within each pairwise comparison. The Entrez Gene identifiers of the human homologs of the genes interrogated by the array were ranked according to the moderated t-statistic computed between the two experimental conditions. Mouse genes with multiple human homologs (or vice versa) were removed prior to ranking. The ranked list represented only those human genes that matched exactly one mouse gene. This ranked list was then used to perform preranked GSEA analyses (default parameters with random seed 1234) using the Entrez Gene versions of the Hallmark, Biocarta, KEGG, Reactome, Gene Ontology (GO), and transcription factor and microRNA motif gene sets obtained from the Molecular Signatures Database (MSigDB), version 5.0.^41^

The overlap between the different nodes was visualized by the generation of an enrichment map by using the results from the GSEA analysis which were input into the Cytoscope software suite (version 3.5.1).^42^ The map was constructed with a cutoff of p < 0.05 and an FDR cutoff of q < 0.25. The similarity cutoff was 0.5 with a combined constant of 0.5. The MCL cluster algorithm was used to group the nodes into clusters.

### Quantification of [Ca^2+^]_i_ oscillation drift

Data was analyzed in GraphPad Prism 7 software (La Jolla, CA) that allowed for interpolation of data points. The first peak after a stimulus was disregarded. The beginning of an oscillation period was determined as the time point prior to the start of a peak rise (t_start_). The period (p) was defined as the difference in time between two adjacent start times (p = t_start+1_ - t_start_). The periods were then plotted as a function of elapsed experimental time and a linear regression was fit to the data. The absolute value of the slope of the linear regression (Δp / time) to this plot was defined as the drift. For determining drift from literature data, insulin or [Ca^2+^]_i_ traces were imported into Adobe Illustrator CS5 (San Jose, CA) and the period measured as described above using the scale bars in the figures. The data analyzed corresponded to references 7,8,12,13,16,18, 43-47, for the *in vitro* data and 5,6,14,48-56 for the *in vivo* data. For testing statistical significance, unless otherwise stated, a paired, twotailed t-test was employed.

## Results and discussion

As mentioned previously, if synchronization produces highly stable [Ca^2+^]_i_ oscillation periods, unique gene expression profiles may then be produced. In the experiments described below, synchronization was achieved using a mimic of classic insulin-glucose feedback loops where a glucose wave (11 +/- 1 mM) with a 5-min period was delivered to a population of islets. In the control experiment, another population of islets was given the same dose of glucose, but without the oscillation. The stability of the [Ca^2^+]_i_ oscillations were then compared, and the gene expression profiles, as determined by microarrays, were evaluated.

## Comparison of [Ca^2+^]_i_ oscillations in synchronized and non-synchronized islets

When a population of islets were exposed to a 5-min glucose wave, 93.5% (58/62) of islets showed one oscillation mode (“slow” oscillations having a period between 3-8 min), which persisted throughout the duration of the experiment. These slow oscillations are what would be expected for production of the 5-min pulses of insulin that are observed *in vivo.* In the constant glucose experiments, subpopulations of islets showed “fast” (oscillation period < 2 min, 9/62, 14.5%), “slow” (29/62, 46.8%), or “compound” (a combination of fast and slow, 15/62, 24.2%) oscillation modes. These variable modes have also been observed previously in other reports.^57, 58^ In addition, several islets in the constant glucose experiment displayed oscillations that started or stopped at different times during the experiment (all data is available in the Supplementary Information and for a summary see S1 Fig and S1 Table). Overall, these data indicated that the islets exposed to the glucose wave had a high homogeneity in their [Ca^2+^]_i_ response, whereas the islets exposed to the constant glucose dose were highly heterogeneous. (Fig 1).

**Fig 1.**
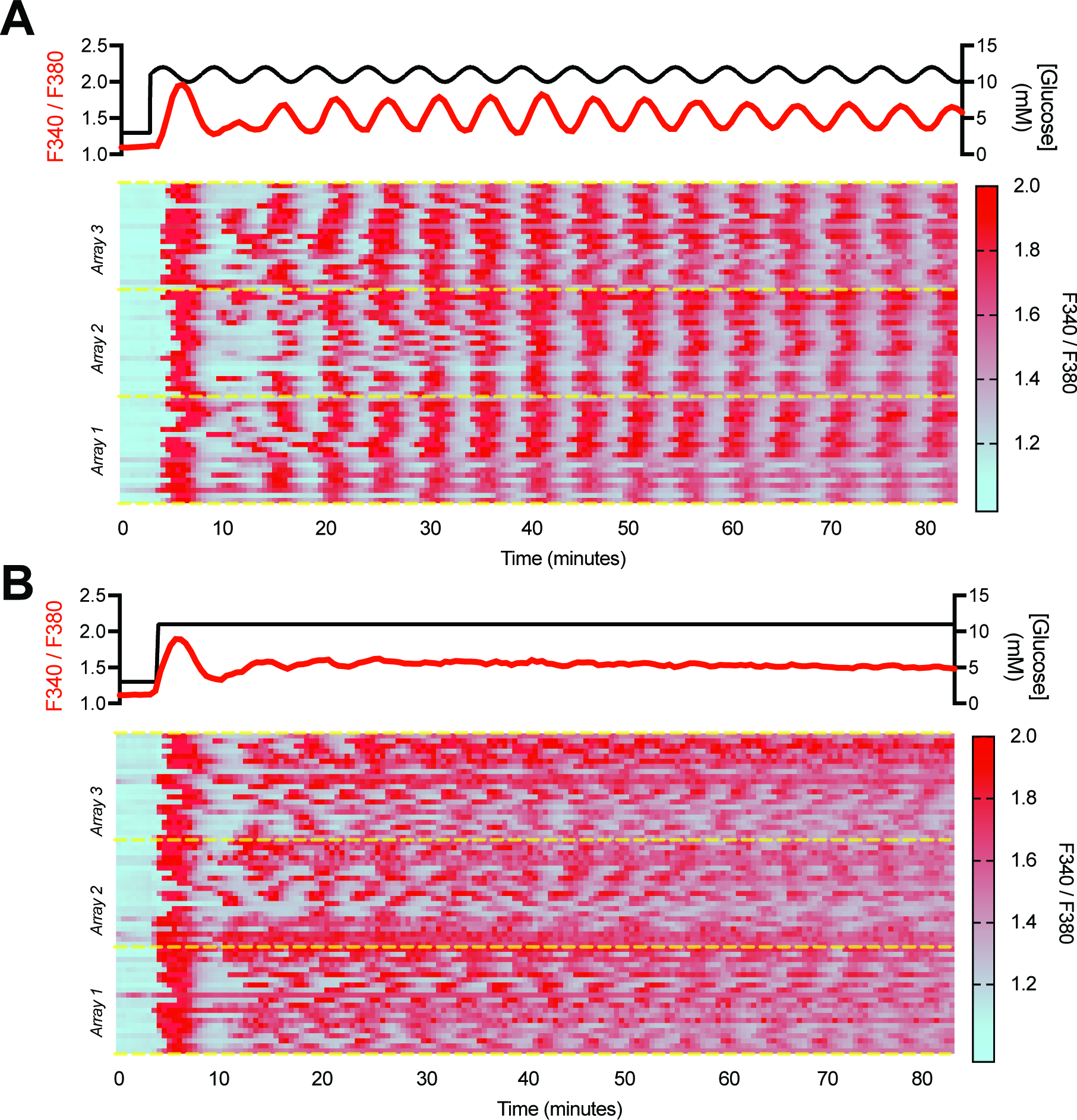
Effect of glucose profiles on islet response. The administered glucose dose (black trace) is plotted on top of each panel. The average F340/F380 signal from all islets in each glucose protocol is plotted at the top of the panels in red. The individual islet signatures are shown in a heatmap. The groupings for the microarrays are delineated with yellow dotted lines. A. When exposed to 11 +/- 1 mM glucose wave with a 5-minute period, the islets synchronized as seen by the homogeneous F340/F380 ratio across the population. B. When exposed to a constant 11 mM glucose signature, the islets oscillated at unique periods with different phases to one another resulting in an incoherent population.

The [Ca^2+^]_i_ oscillations were further quantified to determine the differences in the populations. Both sets of islets had a statistically similar average oscillation period (4.94 vs. 5.10 min for the wave and constant stimulations, respectively), but the standard deviation was 40% lower for the synchronized islets (0.6 min) as compared to the non-synchronized population (1.0 min), indicating a smaller distribution of oscillation periods in the former case. As further demonstration of the consistency in the [Ca^2+^]_i_ periods, the traces corresponding to the grouping for each microarray were averaged and spectral analysis was performed. When synchronized, the phase of the islets relative to the glucose wave was regular, resulting in a conserved [Ca^2+^]_i_ signature after averaging and a pronounced frequency at 0.2 min^-1^ (Fig 2Ai). In contrast, when this was performed with the islets exposed to the constant glucose level, no prominent oscillation frequency was observed (Fig 2Aii) after averaging because the [Ca^2+^]_i_ oscillations had different frequencies, phase offsets, or drifts which resulted in an elevated, but relatively flat average [Ca^2+^]_i_. To illustrate the drift in the [Ca^2+^]_i_ oscillations, the period of successive peaks were plotted for representative [Ca^2+^]_i_ traces for the two glucose treatments (Figs 2Bi and 2Bii). When exposed to the glucose wave, islets entrained to the driving frequency and maintained a constant period (Fig 2Bi). Conversely, when exposed to constant glucose, the period of [Ca^2+^]_i_ oscillations in this representative islet drifted from 8.6 to 4.5 min (Fig 2Bii). The presence of drift was inspected in all islets showing “slow” oscillation modes exposed to the wave (n=58/62) and constant (n = 29/62) glucose treatments. The average drift was significantly lower for islets exposed to the glucose wave (0.12 ± 0.01 minutes / minute) compared to those exposed to the constant glucose level (0.23 ± 0.02 minutes / minute) (p = 0.001). (Fig 2C). In summary, the delivery of an oscillatory glucose level as a mimic of an *in vivo* feedback loop produced a homogeneous [Ca^2+^]_i_ signature from the islet population, which reflects the expected *in vivo* population behavior. Conversely, islets exposed to a constant glucose level produced incoherent signatures of [Ca^2+^]_i_ and would not produce the expected *in vivo* population behavior.

**Fig 2.**
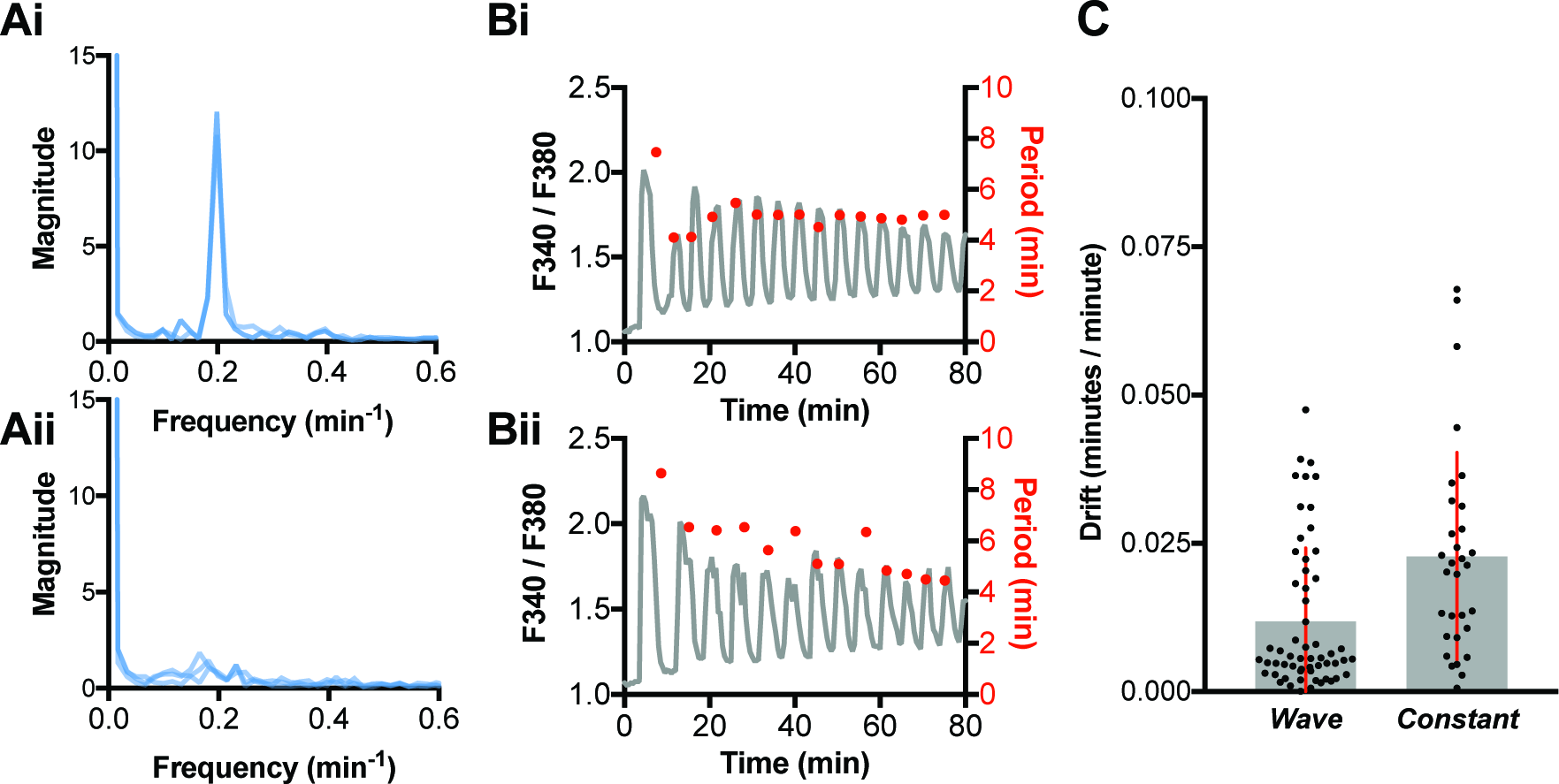
Analysis of islet calcium signatures. **A.** The average spectral analysis for the wave and constant glucose treatments from Figure 1 A and B are shown in Ai and Aii, respectively. B. Representative Fura-2 traces (gray lines) from islets exposed to wave and constant glucose treatments are shown in Bi and Bii, respectively. The red circles above the traces are the measured period for each oscillation corresponding to the right y-axis. C. The average drift measured from islets showing a “slow” oscillation mode is given by the gray bars with the error bars corresponding to ± 1 SD. Individual values of the drift are plotted as black circles (•).

## In Vivo pulsatility is more regular than in vitro

Because other mechanisms for synchronizing islets have been proposed, we hypothesized that we could complement our findings by examining *in vivo* rhythms from published reports. We expected that data obtained *in vivo* would contain less drift than data obtained *in vitro* due to the presence of any synchronizing factor (glucose-insulin feedback or any other mechanism) in the former case. In addition, this comparison served to test the validity of the drift observed in the experiments conducted in our study.

Data were examined from 17 *in vivo* insulin profiles from different species (dogs, rats, mice, humans) across 12 published studies and compared to 32 *in vitro* insulin or [Ca^2+^]_i_ profiles from different species (rat, mice) from 11 published studies. Drifts in the measured periods from representative *in vitro* and *in vivo* experiments are shown in Figs 3A and 3B, respectively. The drift in insulin oscillation periods *in vivo* was significantly lower compared to those measured *in vitro* (0.02 ± 0.01 minutes / minute vs. 0.06 ± 0.05 minutes / minute, respectively, p = 0.011) (Fig 3C). Although these data cannot be considered comprehensive, the results suggest that the presence of a synchronizing factor *in vivo* may help to maintain a constant insulin oscillation period, and therefore [Ca^2+^]_i_ period, similar to what was observed in the population of islets exposed to the oscillatory glucose stimulation (Fig 1A).

**Fig 3.**
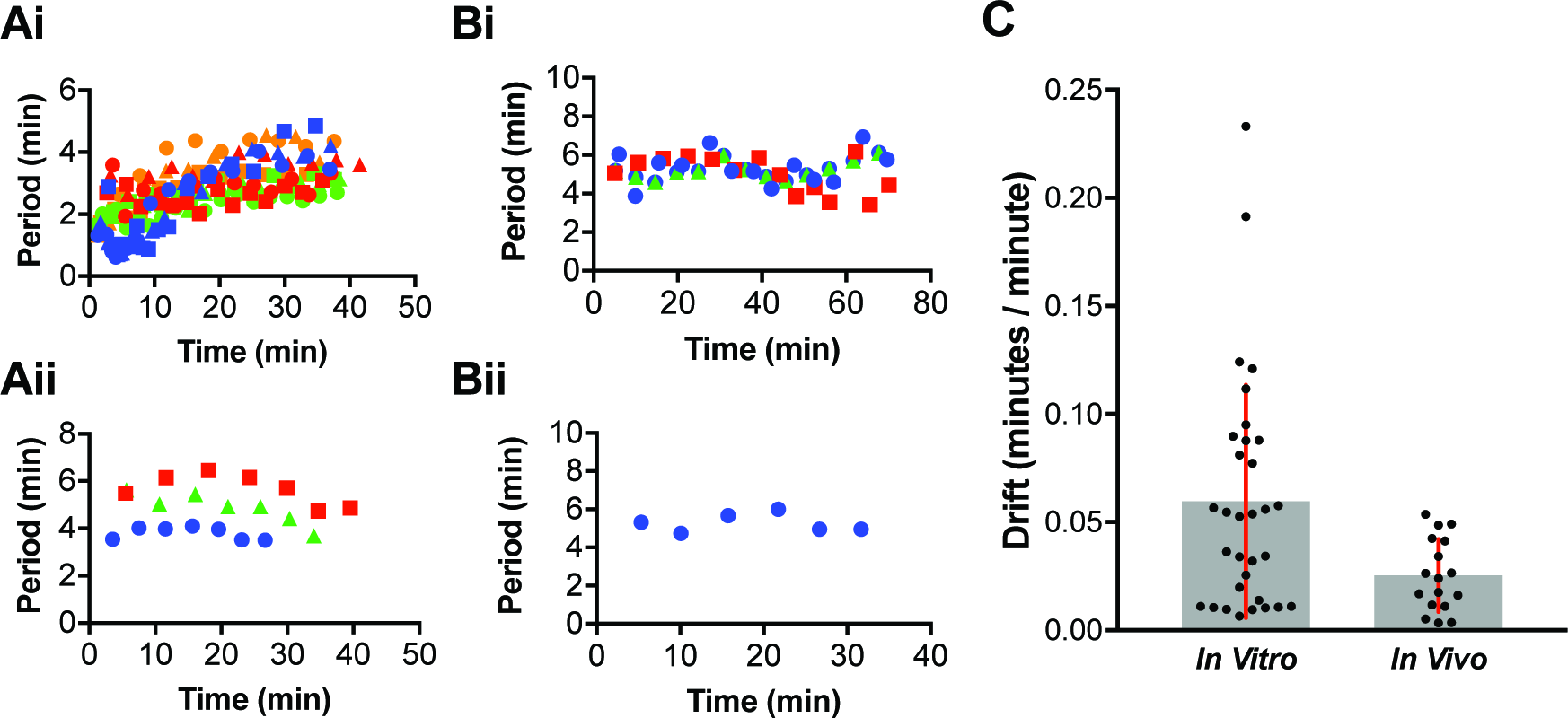
Comparison of drift from in vivo and in vitro data. (A) Shown in Ai and Aii are examples from references 16 and 44, respectively, of the period of [Ca^2+^]_i_ oscillations *in vitro* as a function of experimental time. The various colors and symbols represent different experiments. (B) Shown in Bi and Bii are examples from references 4 and 53, respectively, of the period of insulin oscillations *in vivo* as a function of experimental time. (C) The drift from each experiment *in vitro* and *in vivo* are plotted as black circles (•). The average is shown as a gray bar with the error bars corresponding to ± 1 SD. The values are statistically different (p = 0.011).

Similar results were observed ~25 years ago when *in vitro* hormone pulsatility in isolated islets was previously shown to be more irregular when compared to *in vivo* measurements in the rhesus monkey.^59^ The phase relationships of insulin, glucagon and somatostatin that were observed *in vivo* were lost *in vitro.* In addition, the amplitude and frequency were statistically higher and faster *in vitro* compared to *in vivo.* However, the drift in the oscillations was not examined.

## Differential expression of genes in synchronized islet populations

In biological systems, Ca^2+^ modulation offers a mechanism for cells to implement specificity and efficiency to cellular signaling. Cells have been shown to produce specific responses as a function of [Ca^2+^]_i_ oscillation frequency.^24, 28^ Moreover, in some cases gene expression was found to be regulated with higher sensitivity as a function of the frequency of the oscillatory [Ca^2+^]_i_.^27, 29^ We therefore hypothesized that islets with a consistent [Ca^2+^]_i_ oscillation period may produce specific gene expression compared to islets with a heterogeneous and drifting period, and these differential genes may result in an advantage for those synchronized islets. This advantage may be in the form of more efficient utilization of both tangible (e.g., nucleotides, amino acids, nutrients, ribosomes, energy) and/or intangible (e.g., time)^60^ cellular resources. This may allow islets in a synchronized state to respond and adapt to elevated glucose levels more efficiently, and with less resource consumption than non-synchronized islets.

To test this hypothesis, mRNA levels from islets exposed to the two glucose stimulation protocols were determined using microarray analysis. Initial tests on the array results were performed by principal component analysis (Fig 4A). The first component accounted for 25% of the variability and demonstrated well-defined separation. Clustering along the second component was low, but we attribute this to both islet populations having similar responses to the same dose of glucose, with the only difference being the drift in oscillation period. The larger proportion of low p-values compared to high p-values (Fig 4B) confirmed differential expression of genes between synchronized and non-synchronized islet populations. 400 genes were deferentially expressed with p-values < 0.01 and over 2,000 with p-value < 0.05 (Fig 4C). The number of differential genes expressed between stimulation protocols is not large, but considering that the glucose dose was the same for both sets of islets and that both sets induced oscillations of [Ca^2+^]_i_, the finding of differentially expressed genes between the two treatments, emphasizes the significance of a drifting oscillation period.

**Fig 4.**
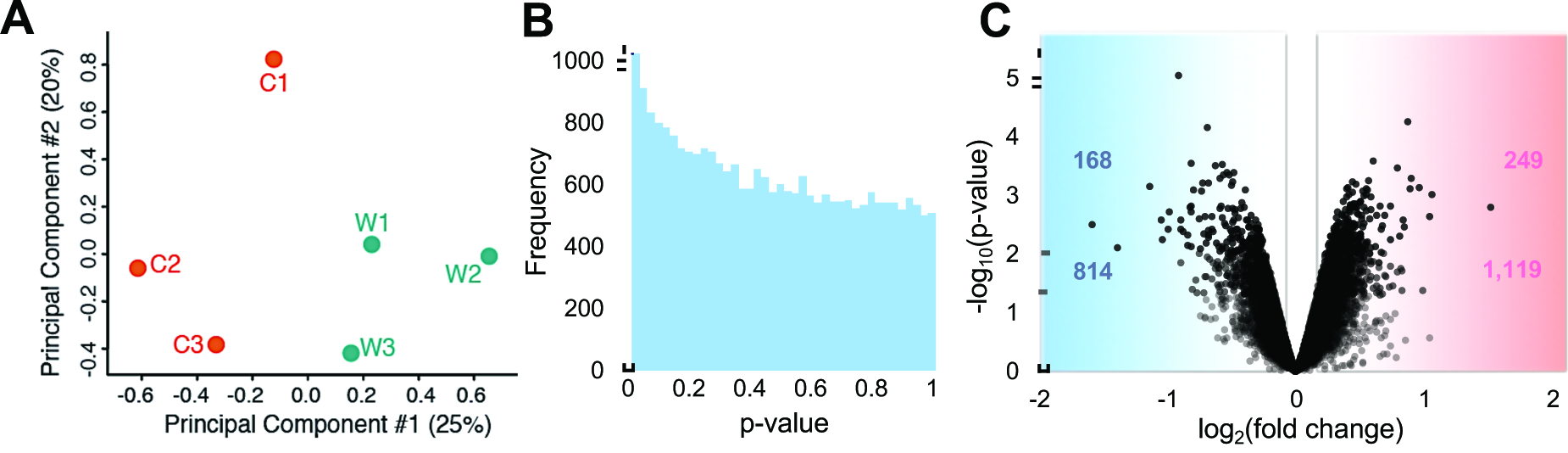
Overview of microarray statistics. (A) Principal component analysis showing separation along the first component. The replicates for the constant glucose treatment are plotted as orange circles and labeled C1-3. The replicates for the glucose wave treatment are plotted as teal circles and labeled W1-3. (B) Histogram of p-values (bin size = 0.001) for the differential comparison. (C) Volcano plot of the microarray results shown as the negative log_10_ of the adjusted p-value against the log_2_ of the fold change. Each circle corresponds to a single gene. Genes up-regulated under the glucose wave are shown as a positive fold-change (red) and the down-regulated as a negative fold-change (blue). Horizontal dashed lines are shown for p = 0.05 and 0.01 thresholds. The number of genes above these thresholds are annotated for the down and up-regulated genes in blue and pink, respectively.

Differential genes were filtered to p-values < 0.01, fold-change > 1.3, and an average RLE signal > 4, producing 68 down-regulated and 23 up-regulated genes. A selection of these genes is shown in Fig 5. The majority were protein-coding with several Ca^2+^-dependent genes down-regulated in the synchronized islets (Camkk1, Calb1, Cdc109b, Pcdha10, Pla2g7). Calmodulin dependent kinase (Camkk1) and Calbindin 1 (Calb1) have been shown to be involved in the calcium/calmodulin-dependent (CaM) kinase cascade that has been implicated in decoding properties of Ca^2+^ dynamics in several cell types.^61^ In islets, CaM-related pathways have been shown to activate insulin expression.^61^ In addition to the antioxidant, glutathione peroxidase 7 (Gpx7) and the serine protease inhibitor 1 (Sepina1b), several transcription factors (Klf6, Klf8, Zfp119a, Zfp46, Fgf14) were also found to be down-regulated.

**Fig 5.**
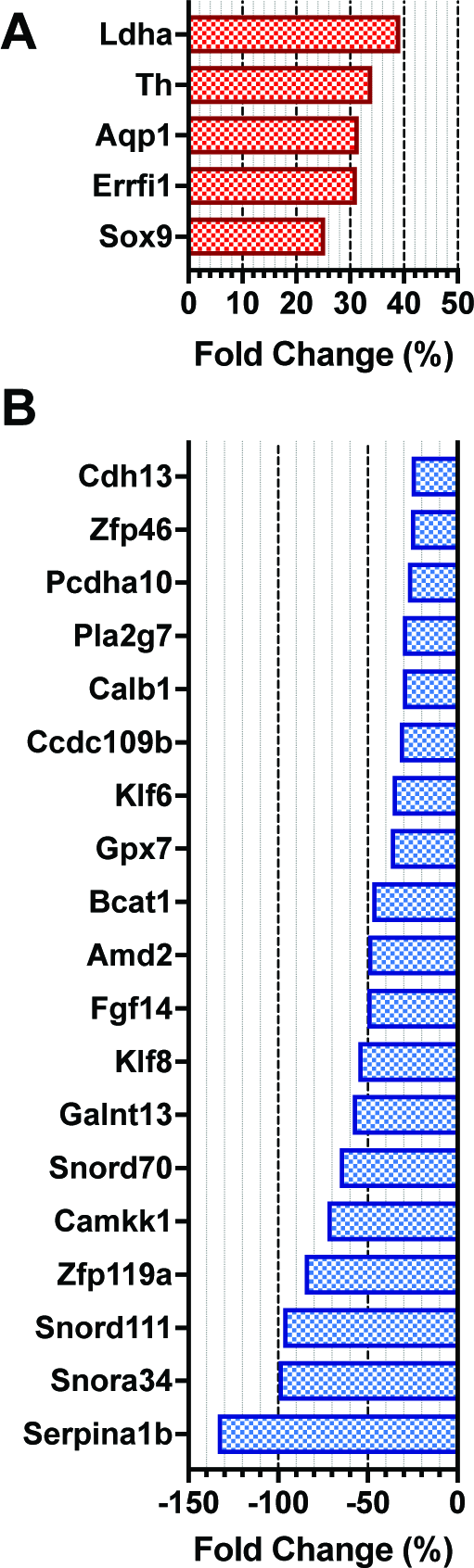
Partial list of differentially regulated genes. Shown is a partial list of differential genes that were up (A) or down (B) regulated in the glucose wave vs. constant treatment. The fold-change is presented as a percentage above or below the fold-change in the constant glucose treatment.

Kruppel-like transcription factors (Klf) have previously been identified as regulators of metabolism and insulin biosynthesis.^62^ Zinc finger proteins (Zfp) have also been connected with islet function and alpha cell identity.^63^ Only 16 genes (23.5%) were found to be pseudogenes while 6 (8.8%) were either small nuclear RNAs (Snor) or long non-coding RNAs (lncRNA).

From the up-regulated transcription factors, Sox9 has been previously identified as a central regulator of pancreogenesis and homeostasis.^63^ Lactate dehydrogenase (Ldha), which is involved in the conversion of pyruvate to lactate, was up-regulated.^64, 65^ Previously, Ldha levels in islets were found to sensitize insulin secretion.^64^ While generally repressed in islets, perhaps this circumstantial up-regulation is transient and serves to facilitate faster response to the administered glucose challenge. The expression of tyrosine hydroxylase (Th), responsible for the enzyme involved in producing the dopamine precursor, L-DOPA, was increased in the synchronized islets. Aquaporin 1 (Aqp1) has been implicated in secretory granule biogenesis and was also up-regulated in this islet population.^66^ The up-regulated genes contained only 1 (5.3%) pseudogene and 3 (13%) lncRNA or Snor.

Chronically elevated [Ca^2+^]_i_ can result in excitotoxicity.^67, 68^ In a recent study, pancreatic beta cells containing a K_ATP_ channel protein knocked out (Abcc8^-/-^) resulted in chronic membrane depolarization and an elevated level of [Ca^2+^]_i_.^68^ The Abcc8^-/-^ model demonstrated beta cell failure and de-differentiation with calmodulin dependent kinases (Camk1d, Camkk1, Camkk2) and EF-hand Ca^2+^ binding proteins (S100a1, S100a3, S100a4, S100a6, S100a13) up-regulated. In contrast, in the synchronized islet population, calmodulin dependent kinases (Camkk1) and EF-hand Ca^2+^ binding proteins (S100z, S100a8, Necab2, Efcab2) were downregulated. While the S100a4 and S100a6 genes were identified as biomarkers for excitotoxicity in the Abcc8^-/-^ model, these were not part of the gene panel in the arrays we used. The Abcc8^-/-^ model also showed cell-adhesion molecule (CAM) related genes were down-regulated. In the synchronized islet population, cadherin13 (Cdh13) was down-regulated, but the CAM gene set was up-regulated with 49 of the 114 genes associated with this KEGG database set enriched (Fig 6). In another recent study, disrupted Ca^2+^ handling and homeostasis, along with ER-stress, revealed an upregulation of calpain in pancreatic beta cells.^69^ Calpain is a Ca^2+^-sensitive protease and hyperactivation has been associated with activation of apoptotic pathways and cell death. In the synchronized islet population, calpain6 was found to be significantly down-regulated.

**Fig 6.**
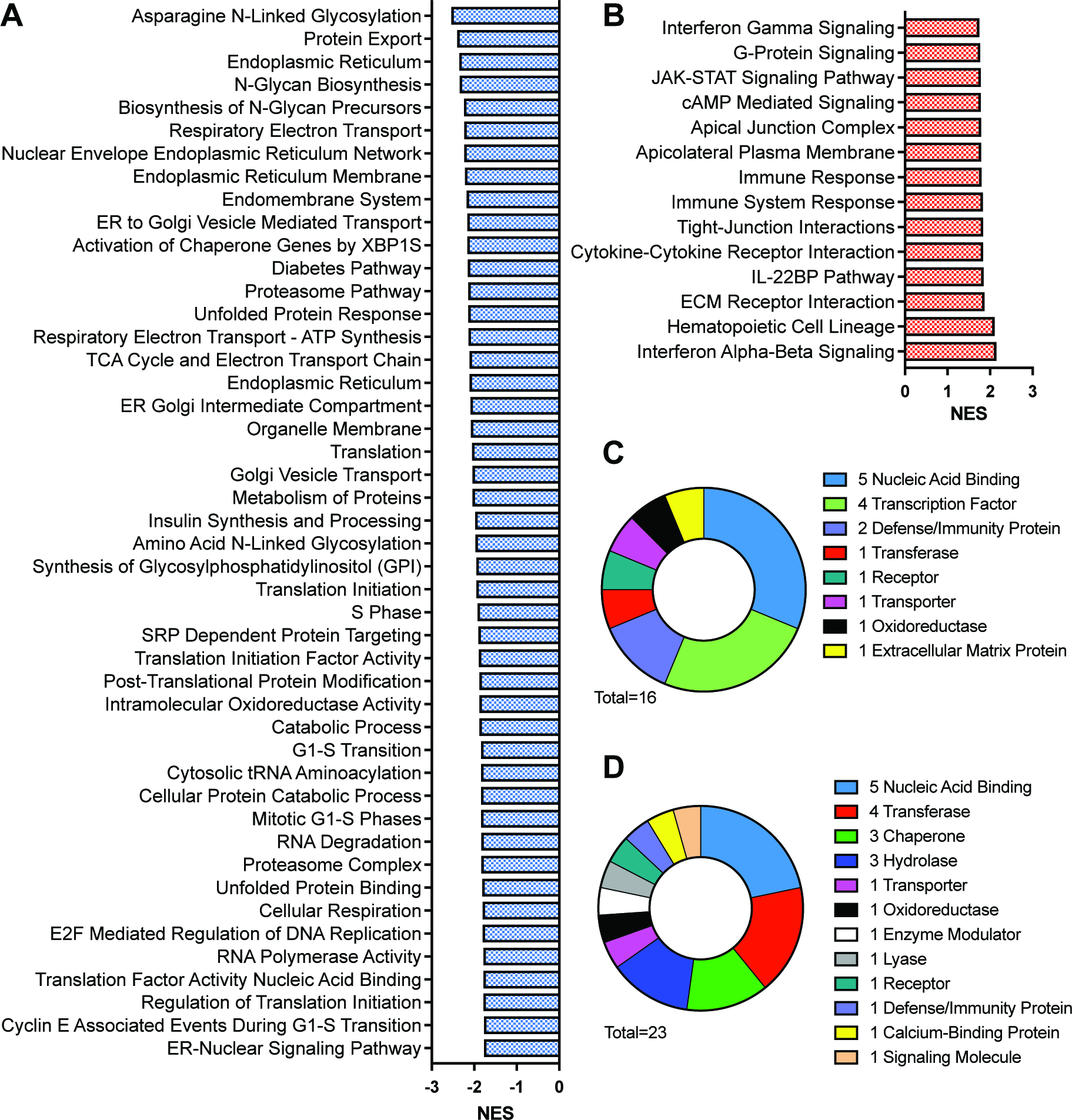
Overview of GSEA and GO results. Gene sets (A) down-regulated and (B) up-regulated in synchronized islets. Only sets with NES > 1.75 are shown, with the full list available in the Supplementary Information. GO results of the protein classes are shown for (C) up‐ and (D) down-regulated genes.

These observations potentially suggest that the synchronized population is more protective, perhaps as a result of less Ca^2+^-excitotoxicity. This better Ca^2+^ handling may occur in the synchronized population because a larger percentage of this group (93.5%) showed slow oscillations of [Ca^2+^]_i_ compared to those under the constant glucose stimulation (50%) (S1 Table). The slow [Ca^2+^]_i_ oscillations may alleviate potential Ca^2+^-dependent cytotoxicity that would occur with fast or continuously elevated [Ca^2+^]_i_. Why the percentage of slow oscillators increased is unknown, but it may have been a result of “recruiting” those islets that showed fast or elevated [Ca^2+^]_i_ oscillations to slow oscillators, as has been observed in a previous report when oscillatory glucose levels were delivered to islets.^10^

To further compare the two populations of islets, GSEA analysis was used to examine differentially regulated groups of genes. This type of analysis has been shown to be useful in identifying diabetic retinopathy^70^, mature onset diabetes of the young (MODY)^71^, type I and II diabetes^40^, and cancer subtypes^72^ amongst others. The gene sets were filtered to normalized enrichment scores (NES) > 1.7 and FDR q < 0.25, which produced 59 sets that were up-regulated and 286 that were down-regulated. The complete list of gene sets is presented in Fig 6 and an enrichment map is given in Fig 7.

**Fig 7.**
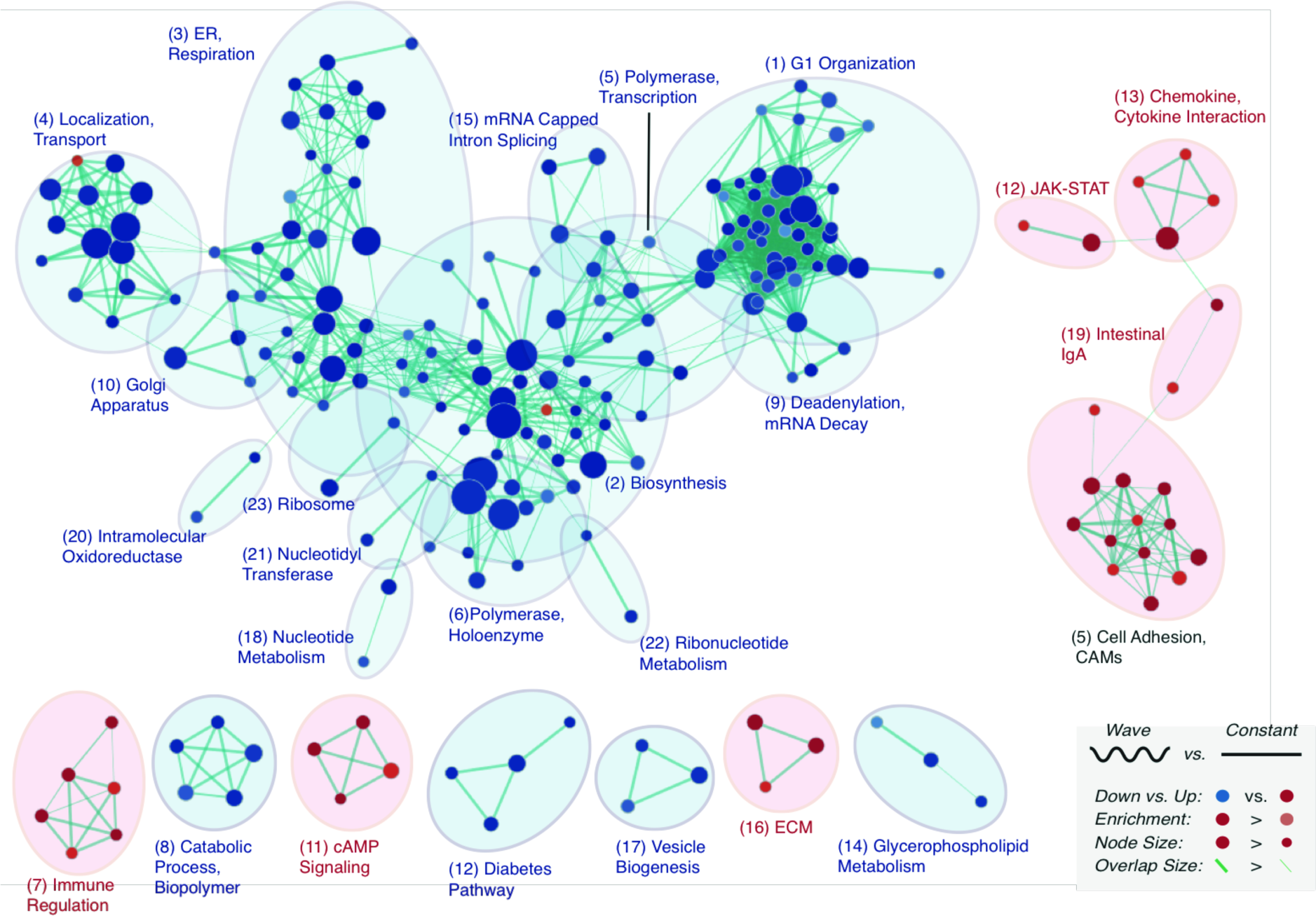
Enrichment map analysis. An enrichment map was created to visualize the general gene expression landscape. A key is shown in the bottom right of the figure. Each node corresponds to a gene set and the thickness of the lines connecting the dots is proportional to the overlap. Genes up-regulated in the wave treatment are red and those down-regulated are blue. Clustering is visualized with circles over the nodes. The clusters are ranked by the number of common nodes and the rank is shown in parentheses.

Overall, there appears to be a general decrease in protein turnover in the synchronized population as indicated by down-regulation of the gene sets for protein export, proteasome pathway, endoplasmic reticulum (ER)-related transport, and unfolded protein response (UPR). UPR is a process that is induced to alleviate proteotoxic stress due to high levels of protein flux to the ER.^73^ Excessive activation of the UPR, and an inability to cope with ER stress, can result in the induction of apoptotic signaling pathways.^74, 75^ More specific to islets, UPR pathways induced by high glucose have been implicated in beta-cell dysfunction and death.^74^ Furthermore, the ER has well-documented Ca^2+^ stores, making it a plausible target for the action of finely modulated [Ca^2+^]_i_ oscillations. In general, the down-regulated gene sets suggest a quiescent-like state in the synchronized islets. Terms related to protein expression and modification (proteasome pathway, translation, metabolism of proteins, and translational initiation) were all down-regulated. Protein synthesis is one of the most energy-demanding functions in cells^76^ so the ability to down regulate these processes, even under a high glucose load, would likely enable better allocation of resources to other cellular functions in an economic and efficient manner. Catabolic processes were also down-regulated in synchronized islets, which is intriguing in these elevated glucose states as catabolic pathways have been found to be activated in times of nutrient stress to fuel and provide building blocks for other cellular processes.^77,78^ Potentially, the synchronized islets had a more efficient response to the elevated glucose, leading to fewer intracellular resources being used in response-
and adaptation-related pathways, compared to the non-synchronized islets.

In addition to reduced protein translation, insulin synthesis and processing was found to be down-regulated in the synchronized group. The reason for lowered synthesis and processing of insulin may be related to a more efficient use of the hormone. *In vivo*, pulsatile insulin more effectively lowers elevated glucose levels compared to a constant administration at the same dose.^30^ This implies that the action of an insulin molecule secreted in pulses and in phase with other islets, is more efficient than an insulin molecule secreted from an islet that is not synchronized. Therefore, since the insulin is more efficient, less insulin can be produced in these islets, which allows utilization of those intracellular resources for other functions.

The synchronized population also had down-regulation of the electron transport chain (ETC)-, TCA cycle‐ and respiration-related gene sets. In addition to these pathways being involved in energy production, the ETC is a primary source of electron leakage and the production of reactive oxidative species (ROS). Islets are known to have low levels of antioxidant enzymes and are susceptible to damage from elevated levels of ROS [79]. As a result, down-regulation of the ETC-related pathways may be protective by mitigating potential ROS production. Several other down-regulated gene sets in the synchronized population may also minimize the production of detrimental byproducts. For example, high protein turnover and UPR-associated pathways were down-regulated, and both have been affiliated with fibril and plaque formation in islets, resulting in failure and death.^75^ Similarly, the formation of glycosaminoglycans has been connected with diabetic pathogenesis and was found to be down-regulated.^80^

Finally, there appears to be higher control of pathways related to cell morphology. Gene sets for CAM, ECM-receptor and tight-junction interactions were up-regulated in the synchronized islet population. This is in direct contradiction to the Abcc8^-/-^ islets which had chronically elevated [Ca^2+^]_i_ levels and showed downregulated cell adhesion molecules. Furthermore, independent of the Abcc8^-/-^ islets, in normal islets, excitotoxicity has been shown to be detrimental to islet morphology and structure^68,81,82^; therefore, the recruitment of non-oscillating islets by a glucose wave would induce a protective function.

In other cell types, [Ca^2+^]_i_ oscillations were found to effect the expression of transcription factors and amplify signaling pathways. In our findings, several signaling pathways were also found the be up-regulated in the synchronized group, notably, cAMP mediated signaling. This pathway has been shown to be involved in amplification of glucose-induced insulin secretion.^83^ When exposed to elevated glucose, single beta cells and isolated islets were shown to produce cAMP oscillations with similar periodicity and phase as oscillations of [Ca^2+^]_i_.^84^ Furthermore, increases in [Ca^2+^]_i_ were postulated to magnify the cAMP signatures. As a result of this relationship between cAMP and [Ca^2+^]_i_, homogeneous and consistent [Ca^2+^]_i_ profiles observed in synchronized islets would also be expected to result in similar cAMP signatures. Therefore, the cAMP dynamics in this population are expected to result in an efficient and programmed cellular response. Despite cAMP being connected with insulin secretion, the insulin synthesis pathway was down-regulated in this population.

Finally, the “diabetes pathway” which contains a list of 136 genes in the Reactome database, was found to be down-regulated in the synchronized islet population (68 genes were in the leading edge with an NES of ‐2.14 and a q-value of 0.0015). The majority of the genes in the set are associated with binding (47.3%) and catalytic activity (38.2%). The down-regulation of the diabetes pathway is attributed to the low bioenergetic state in the synchronized population as low protein turnover would impose low proteotoxic stress and may alleviate high energy requirements. Further experiments are required to fully characterize and test this hypothesis.

## Conclusion

In this report, *in vivo* islet synchronization was mimicked by delivering a low amplitude glucose wave. While there may be additional or alternative routes to islet synchronization *in vivo*, the result would nevertheless be a synchronized population of islets that would have coincident oscillations of [Ca^2+^]_i_. It was found that a synchronized population had a lower drift in Ca^2+^-oscillation periods compared to a non-synchronized population. The synchronized islets showed reduced utilization of intracellular resources as expression of genes related to protein turnover, protein translation, and energy production were lowered, while those that were related to maintenance of cellular morphology were increased. Combined, these results appear to be in agreement with the hypothesis that a synchronized islet population, like those observed in healthy individuals, would be able to respond more efficiently, with respect to intracellular resources, to elevated glucose levels.

## Acknowledgements

This work was supported by a Planning Grant from Florida State University. NM was supported by an AHA Predoctoral Fellowship. Microarray analysis was performed at the Boston University Microarray and Sequencing Resource and the Boston University Clinical and Translational Science Institute (CTSI), which is supported by a CTSA grant (1UL1TR001430). The authors would like to also gratefully acknowledge Mr. Basel Bandak for assistance with islet procurement.

## Supporting information

**Fig S1.**
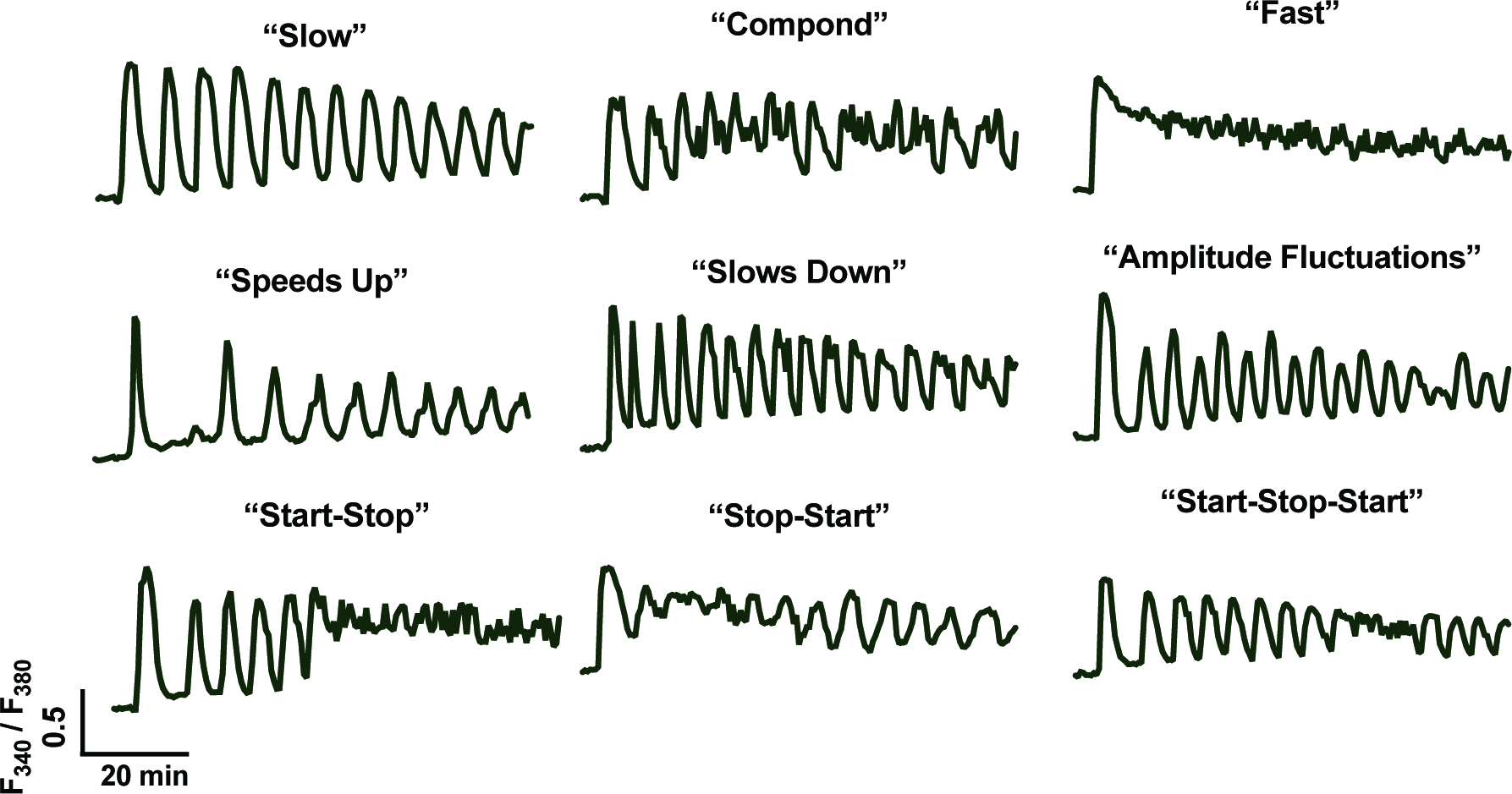
Islet Ca^2+^ oscillation types. Ratiometric fluorescence signal for the Ca^2+^ imaging is plotted as a function of time for eight islets exposed to 11 mM glucose. The Top Row shows different types of oscillators. The Middle Row shows distortions that happen during the span of a regular in respect to period and amplitude. The Bottom Row shows different start and stop behavior.

**Table 1.** Distribution of oscillator types. Distribution of oscillator types in the glucose wave (“Wave”) and the constant glucose (“Constant”).

| Type | Constant | Wave |
| --- | --- | --- |
| Slow Oscillator (%) | 46.8 | 91.9 |
| *Avg (min)* | 5.18 | 5.10 |
| *SD (min)* | 1.16 | 0.66 |
| Compound Oscillator (%) | 24.2 | 3.2 |
| Fast Oscillator (%) | 14.5 | 0.0 |
| Stops Oscillating (%) | 11.3 | 4.8 |
| Not Oscillating (%) | 3.2 | 0.0 |

